# PNGaseL from *Flavobacterium akiainvivens* targets a diverse range of N-glycan structures

**DOI:** 10.1101/2025.05.27.656428

**Authors:** Cassie R. Bakshani, Paulina A. Urbanowicz, Concepcion Badia Tortosa, Javier Mauricio Melo Diaz, Magdalena Kujawska, Taiwo O. Ojuri, Lindsay J. Hall, Daniel I.R. Spencer, David N. Bolam, Lucy I. Crouch

## Abstract

PNGases are used by a wide range of organisms to remove N-glycan structures from proteins for use as either nutrients or in glycoprotein processing. PNGaseF is the most well-characterised enzyme of this family and is widely used in glycobiology to allow study of the N-glycome of a specific protein, cell and tissues, for instance. Despite this, PNGaseF has limitations in the types of N-glycan structures it can target. In this study, we explored the specificities of six uncharacterised PNGases selected from diverse parts of the PNGaseF superfamily. One of these enzymes, PNGaseL from *Flavobacterium akiainvivens*, is the highlight of this study due to its very broad specificity exemplified by its ability to cleave mammalian-, plant- and invertebrate-type complex N-glycans as well as high-mannose N-glycans. A detailed biochemical and structural characterisation was carried out against a variety of substrates to illustrate the advanced capability of PNGaseL in comparison to the canonical PNGaseF and PNGaseA enzymes. To determine the optimal reaction conditions, assess stability and define limitations of PNGaseL, a series of validation studies was performed. The data reveal that PNGaseL has potential utility in a range of glycobiology applications that are superior to the current commercially available options.

## Introduction

N-glycans are common posttranslational modifications to mainly eukaryotic proteins, particularly on secreted proteins, and have a variety of roles including protection from degradation and cell-to-cell communication. A pentasaccharide core (Man3GlcNAc2) is ubiquitous to all eukaryotic N-glycan structures, but elongation of this core structure is categorised into three types: high-mannose, complex and hybrid, which combines features of both other types (figure 1)^1^. High-mannose N-glycans have highly conserved structures across organisms, although the number of mannose sugars per glycan may vary^1^. In contrast, complex N-glycan structures vary significantly between kingdoms and there are characteristic decorations recognised for mammals, plants and invertebrates, for example^1^. ‘N-glycome’ studies have also revealed heterogeneity in terms of the variety of N-glycan structures attached to individual proteins at different sites^2^.

**Figure 1.**
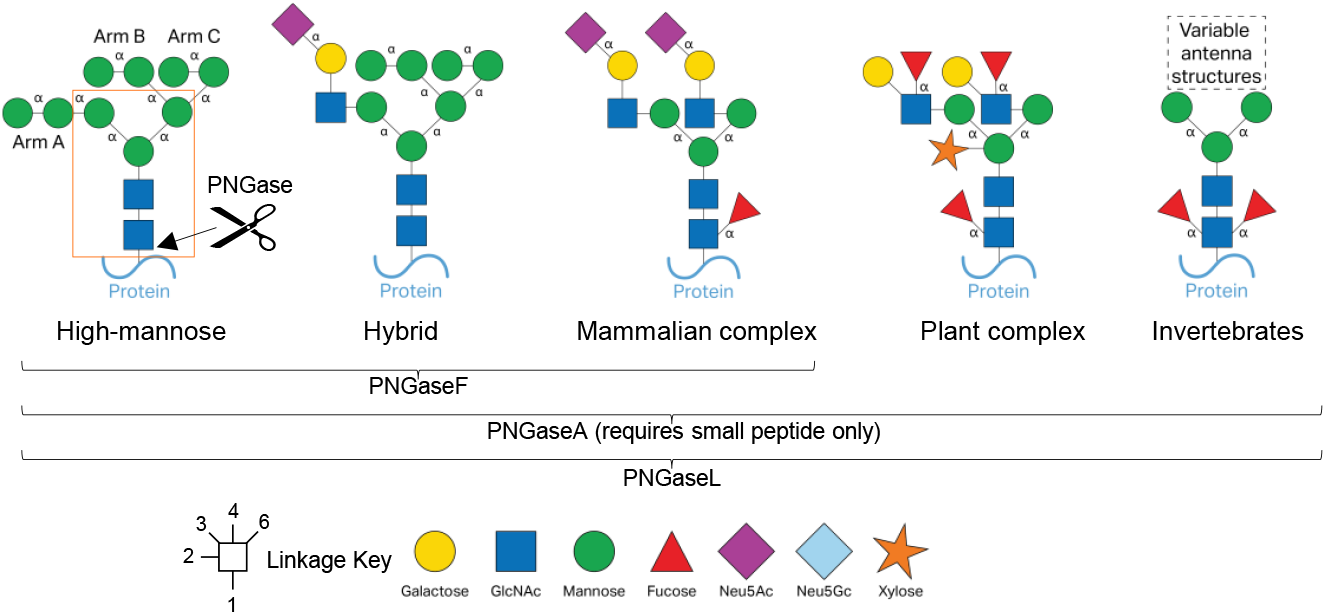
Structures of eukaryotic types of N-glycans. The core of an N-glycans structure is indicated by the orange box and α-linkages are labelled. High-mannose N-glycans are ubiquitous to many organisms (far left) and only comprise of α-linked mannose. Mammalian complex N-glycans (centre) have LacNAc antenna capped with sialic acid. These types of N-glycans are much more variable than other characterised types with the possibility of up to four antenna (two antenna are shown here) and core (α1,6) or antenna fucosylation. Plant complex N-glycans have Lewis A antenna structures, core α1,3-fucose, and bisecting β1,2-xylose. N-glycans from invertebrates can have both α1,3- and α1,6-fucosylation on the core GlcNAc.

One of the main classes of enzymes that target N-glycans are Peptide:N-glycosidases (PNGases) that hydrolyse directly between N-glycan and protein, and several different enzymes have now been described with specificity towards different types of N-glycans and the state of the protein^3-7^. There are currently two major groupings of PNGase enzymes – PNGaseF and PNGaseA, where the “F” and “A” represent *Flavobacterium* and almond, respectively (*Flavobacterium meningoseptica* was later renamed *Elizabethkingia meningoseptica*). PNGaseF and A are broadly associated with prokaryotes and eukaryotes, respectively. In prokaryotes, these enzymes are used to remove N-glycans from proteins to use as a nutrient source and are typically localised to the cell surface^7^. PNGaseA enzymes expressed by eukaryotes usually act intracellularly to recycle N-glycans. The putative PNGaseA enzymes found in prokaryotes are largely from the Actinobacteria and Proteobacteria phyla. Putative PNGaseF enzymes found in eukaryotes are largely associated with marine and aquatic organisms, such as fish, oysters, starfish, sea cucumbers, and algae.

In glycobiology studies, one of the main enzymes used is PNGaseF as it has a broad specificity for mammalian-type N-glycan structures and well-established protocols exist for its application^8^. The specificity of this enzyme, however, is limited to N-glycans without α1,3-core fucose, so it cannot act on plant or invertebrate-type glycans^9^. PNGaseA, conversely, can remove N-glycans with an α1,3-core fucose, but its preference for N-glycans attached to only a short peptide presents another disadvantage^10^. In this study, we characterise six new PNGase enzymes from the PNGase F superfamily, focussing on candidates from microbes that are not associated with being mammalian commensals or pathogens. This includes a PNGase from eukaryotic origin. We demonstrate that one of these, PNGaseL, exhibits very broad specificity and high potential for biotechnological applications.

## Methods

### Bioinformatics analysis and identification of target enzymes

1897 PNGaseF sequences (IPR015197) were downloaded from InterPro^11^. The multiple sequence alignment was completed in MAFFT v7^12^ and trimmed using trimAL v1.5.rev0^13^ with options “-gt 0.5 -cons 50”. The maximum likelihood phylogeny was constructed in IQ-Tree v2.3.2^14^ based on the best-fitting amino acid substitution model WAG+R10. Structural modelling of PNGase enzymes was completed in ColabFold v1.5.2-patch: AlphaFold2 using MMseqs2^15^. Figures were made with PyMOL^16^. The boundaries of modules were determined using the models and information available in InterPro.

### Cloning and recombinant expression

Genes for the mature forms of the six putative PNGases were synthesised with NheI and XhoI restriction sites at the start and the end of the genes, respectively, and were cloned into pET28b. The sequences were codon-optimisation for expression in *E. coli*. Recombinant plasmids were transformed into Tuner cells (Novagen) and plated onto Luria-Bertani (LB) broth containing 50 μg/mL kanamycin and incubated overnight at 37 °C. Cells were then transferred into 1 L of LB medium (in a 2 L flask) and incubated at 37 °C until mid-exponential phase with shaking at 180 rpm. Cultures were then cooled to 16 °C, isopropyl β-D-thiogalactopyranoside (IPTG) was added to a final concentration of 0.2 mM, and the cultures were incubated overnight at 150 rpm and 16 °C in an orbital incubator. These cells were harvested and recombinant His-tagged protein was purified from the cell-free extracts using immobilised metal affinity chromatography (IMAC). The purity and size of the proteins was assessed by sodium dodecyl sulfate-polyacrylamide gel electrophoresis (SDS-PAGE). The SDS-PAGE gels used were precast 4-16 % gradient (Bio-Rad) and stained with Coomassie Brilliant Blue (ReadyBlue® Protein Gel Stain, Sigma-Aldrich) to visualise total protein. The concentrations of recombinant enzymes were determined using absorbance at 280 nm using a Nanodrop and their calculated molar extinction coefficients.

### PNGase specificity screening

The initial screening for activity was carried out against bovine RNaseB (R7884, Sigma-Aldrich), bovine a1acid glycoprotein (G3643, Sigma-Aldrich), bovine fetuin (F2379, Sigma-Aldrich), horseradish peroxidase (77332, Sigma-Aldrich), phospholipase2 from honeybee venom (P9279, Sigma-Aldrich) and γ-globulins from human blood (G4386, Sigma-Aldrich). Assays were carried out in 20 mM 4-morpholinepropanesulfonic acid (MOPS), pH 7.0, at 37°C, with a final glycoprotein concentration of 10 mg/mL, except for PLA2 which was 1 mg/mL. The final enzyme concentration was 5 μM. The SDS-PAGE gels used were precast 4-16 % gradient (Bio-Rad) and either stained with Coomassie Brilliant Blue (ReadyBlue® Protein Gel Stain, Sigma) to visualise total protein or Schiff’s Fuchsin (Pierce Glycoprotein Staining Kit) to visualise just glycoprotein.

### Thin-layer chromatography

9 μL of recombinant enzyme assays were spotted onto silica plates. The plates were resolved in running buffer containing butanol/acetic acid/water in a 2:1:1 and 1:1:1 ratio for high-mannose N-glycan substrates and complex N-glycan substrates, respectively. They were resolved twice, with drying in between, and stained using a diphenylamine-aniline-phosphoric acid stain^17^.

### Three assay conditions with PNGaseF, A, and L

Glycoprotein samples were treated in one of the following ways: 1) denatured for 10 min at 100°C water (“boiling”), 2) denatured for 10 min at 100°C in 0.5% SDS/40 mM dithiothreitol (“boiling and detergent”) or 3) not subjected to any pre-treatment (“untreated”). Glycoproteins were incubated with PNGase L at 37°C at a final concentration of 2 µM. Additionally, samples exposed to 0.5% SDS/40 mM dithiothreitol were digested in the presence of 1% NP-40. Released N-glycans were converted to aldoses by incubating with 0.1% formic acid for 40 minutes at room temperature, filtered through a protein binding plate (Ludger), and dried using a centrifugal evaporator. The plantibody substrate is a nanobody-Fc fusion protein was produced from *Arabidopsis thaliana* seeds.

### Procainamide labelling

Procainamide labelling was performed by reductive amination using a procainamide labelling kit containing sodium cyanoborohydride as a reductant (Ludger). Excess reagents were removed with HILIC SPE purification plates (Ludger). Membrane was conditioned successively with 200 µL of 70% ethanol (vol/vol), 200 µL of deionized (DI) water, and 200 µL acetonitrile. Procainamide-labelled samples were then spotted on the membrane and allowed to adsorb for 15 min. The excess dye was washed with acetonitrile. Labelled *N*-glycans were eluted with 300 µL of DI water.

### LC-FLR-ESI-MS analysis of procainamide labelled glycans

Procainamide-labelled glycans were analysed by LC-FLR-ESI-MS. Here, 25 µL of each sample (prepared in 25:75 water/acetonitrile solution) was injected into a Waters ACQUITY UPLC (ultra-performance liquid chromatography) Glycan BEH Amide column (2.1 × 150 mm, 1.7-µm particle size, 130-Å pore size) at 40 °C on a Dionex Ultimate 3000 UHPLC (ultra-high-performance liquid chromatography) instrument with a fluorescence detector (fluorescence excitation wavelength [λ_ex_] = 310 nm, fluorescence emission wavelength [λ_em_]= 370 nm) attached to a Bruker amaZon speed ETD. Mobile phase A was a 50 mM ammonium formate solution (pH 4.4), and mobile phase B was neat acetonitrile. Analyte separation was accomplished by gradients running at a flow rate of 0.4 mL/min from 76 to 58% mobile phase B over 70 min for honeybee venom phospholipase A2, ovalbumin, bovine α1acid glycoprotein, bovine fetuin, human serum and plantibody N-glycans, from 72 to 60 % mobile phase B over 45 min for IgA N-glycans and from 70 to 62 % mobile phase B over 35 min for human IgG, bovine RNaseB N-glycans and horseradish peroxidase, respectively. The amaZon speed was operated in the positive sensitivity mode using the following settings: source temperature, 180 °C; gas flow, 41 min^™1^; capillary voltage, 4,500 V; ICC target, 200,000; maximum accumulation time, 50.00 ms; rolling average, 2; number of precursor ions selected, 3; scan mode, enhanced resolution; mass range scanned, 400 to 1,700. The LC chromatograms and MS/MS data was viewed in Bruker Compass.

### LC-FLD analysis of procainamide labelled glycans – for IgG, fetuin and horseradish peroxidase (pH optimisation and stability testing)

Procainamide-labelled glycans were analysed by HILIC-LC. Here, 50 µL of each sample (prepared in 28:72 water/acetonitrile solution) was injected into a Waters ACQUITY UPLC (ultra-performance liquid chromatography) Glycan BEH Amide column (2.1 × 150 mm, 1.7-µm particle size, 130-Å pore size) at 40 °C on a Waters ACQUITY UPLC (ultra-high-performance liquid chromatography) H Class instrument with a fluorescence detector (fluorescence excitation wavelength [λex] = 310 nm, fluorescence emission wavelength [λem] = 370 nm). Mobile phase A was a 50 mM ammonium formate solution (pH 4.4), and mobile phase B was neat acetonitrile. Analyte separation was accomplished by gradients running at a flow rate of 0.4 mL/min from 72 to 62% mobile phase B over 35 min for IgG and horseradish peroxidase N-glycans and from 72 to 53% over 60 min for fetuin samples, respectively.

## Results

### Bioinformatics analysis and selections of diverse uncharacterised PNGases

Six putative PNGase enzymes were selected from the PNGase F superfamily guided by phylogenetic analysis (figure 2). The availability of host/origin background information was the initial criteria used to shortlist putative PNGase candidates. From this shortlist, we selected candidates that represented a wide variety of origins and environmental chemistries. These were from: *Draconibacterium orientale* (PNGaseDo), *Crassostrea gigas* (PNGaseCg), *Runella zeae* (PNGaseRz), *Flavobacterium akiainvivens* (PNGaseL), *Labilithrix luteola* (PNGaseLl), and *Deinococcus radiodurans* (PNGaseDr). *Draconibacterium orientale* is a gram-negative bacterium from the phylum Bacteroidota and class Bacteroidia, isolated from a marine sediment sample from Weihai, China^18^. *Crassostrea gigas* (Pacific oyster) is the only eukaryotic enzyme tested in this study. The N-glycan structures from oysters have been shown to have different characteristics to mammalian N-glycans^19^. *Runella zeae* is a gram-negative bacterium from the phylum Bacteroidota and class Cytophagia and was isolated from the stems of maize^20^. *Flavobacterium akiainvivens* is a gram-negative bacterium from the phylum Bacteroidota and class Flavobacteriia. This microbe was isolated from decaying wood of the Hawai’ian plant *Wikstroemia oahuensis* (ākia) and it is also the state microbe of Hawai’i^21^. *Labilithrix luteola* is a gram-negative mesophile from the phylum Pseudomonadota and Class Deltaproteobacteria and isolated from forest soil samples on Yakushima Island, Japan^22^. Lastly, *Deinococcus radiodurans* is a gram-positive polyextremophile from the phylum Deinococcota and class Deinococci. It has been crowned the world’s toughest bacterium and was originally isolated from tinned food that had been sterilised using gamma radiation^23^.

**Figure 2.**
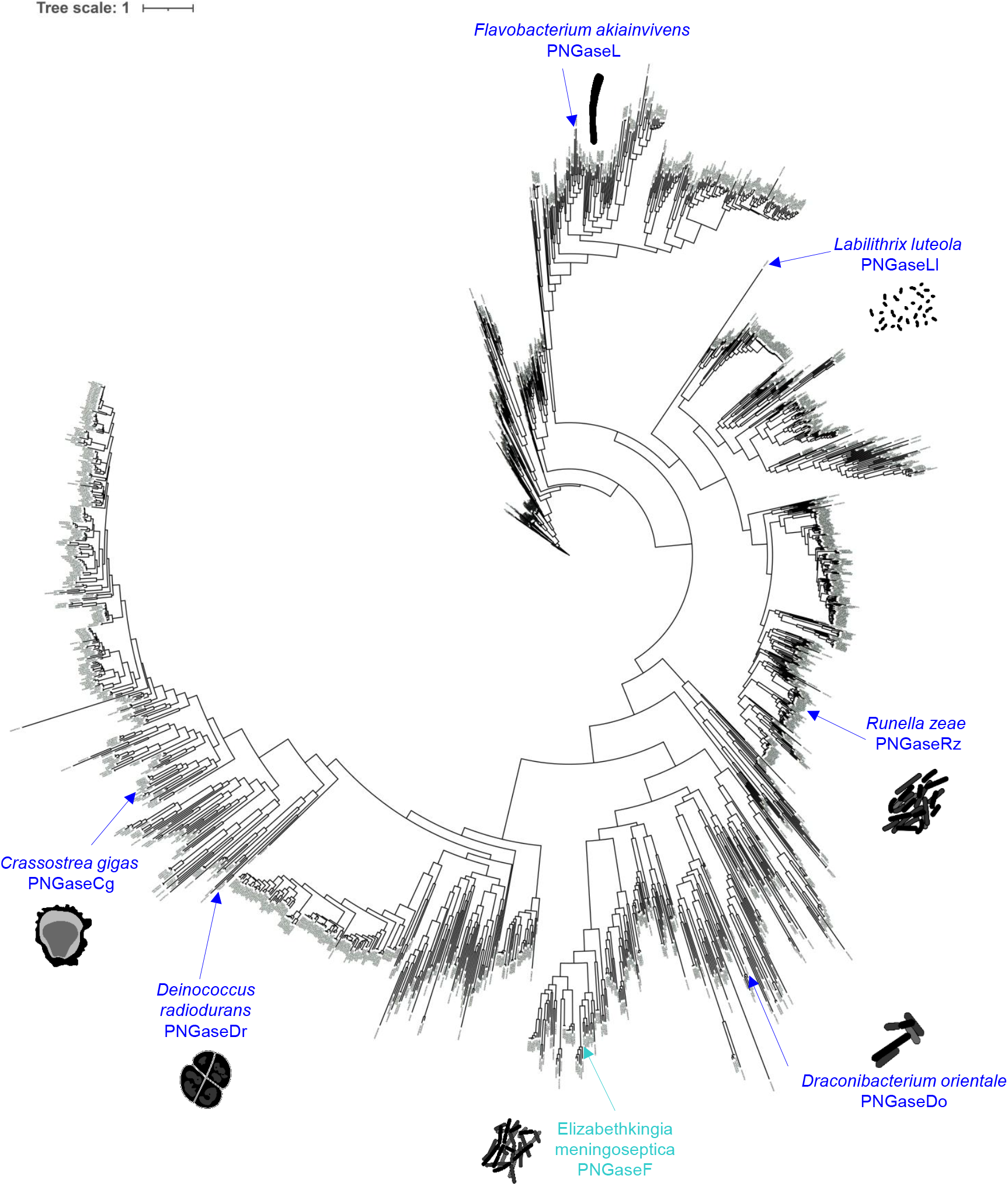
Phylogenetic tree of the PNGaseF superfamily. 1897 PNGaseF sequences were used to construct a phylogenetic tree. The six PNGases characterised in this report and the archetypal PNGaseF are highlighted in dark blue and cyan, respectively. The PNGases were chosen based on being in different clades and the organisms they are derived from existing in different enveronmental niches.

### Structural analysis of selected PNGases

To investigate the predicted tertiary structures of the six PNGase enzymes, we used AlphaFold2 (figure 3). The catalytic modules were all predicted to be two eight-stranded anti-parallel β-sheet structures sitting side-by-side, akin to published PNGase enzyme structures^24^. The accessory modules, however, varied between the different enzymes. PNGaseDo has three β-sheet modules, which likely have linker or carbohydrate binding roles. This arrangement of modules, where the final module is positioned above the active site of the catalytic module has been observed previously, for example a GH20 β-GlcNAc’ase^25^ and a GH29 α-fucosidase^26^. The module positioned over the active site likely has a substrate binding role and siphons substrate towards the active site. PNGaseCg has an α-helix running up the back of the catalytic module (orange), which is connected to a module (“P”) comprised of both α-helices and β-sheets and is a module associated with proteases. P module is also positioned over the active site, so could possibly be binding substrate. PNGaseRz has a β-sheet module that is connected to the catalytic module via a small α-helix. This accessory module is positioned away from the active site and is reminiscent of the N-terminal bowl-like (NBL) accessory module from PNGaseF TypeII from *Elizabethkingia meningoseptica* and B035DRAFT_03341^PNGase^ from *Bacteroides massiliensis*^6,7^. The role of this module is unknown, but in all cases these accessory modules could orient the catalytic module to face outward from the bacterial cell and towards potential substrates. PNGaseL has a small β-sheet module at the C-terminus, which could also possibly interact with substrate. The PNGaseLl structure is a catalytic module and a flexible linker, again likely facilitating the enzyme to project away from the bacterial cell wall. PNGaseDr has a similar modular organisation to PNGaseCg, with an α-helical spine and possible binding modules positioned over the active site. Notably, PNGaseL and PNGaseDo have C-terminal T9SS secretion signals, so these enzymes are likely secreted by their respective organisms.

**Figure 3.**
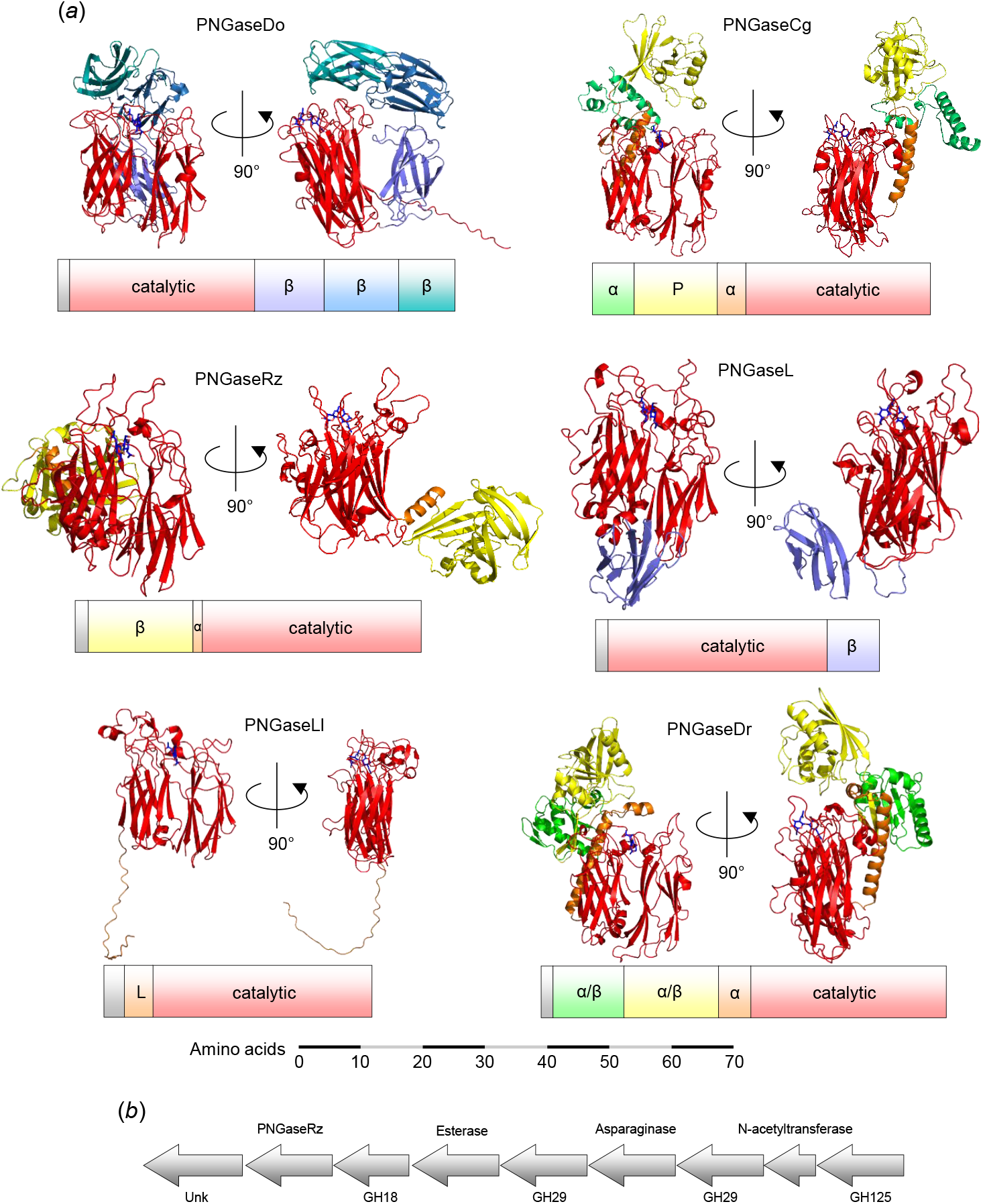
Structures of the PNGase enzymes. (a) Alphafold models of the six PNGase enzymes shown from two different angles with the modular organisation of the polypeptide shown below each model. A scale bar has been included. All the catalytic domains (red) are predicted to be two 8 stranded anti-parallel β-sheets. Modules to the N- and C-terminus of the catalytic domain are coloured lime/yellow/orange and purple/blue/teal, respectively. The signal sequences have not been shown in the structures, but putative linker sequences have been included. The GlcNAcβ1,4GlcNAc (dark blue) from PNaseF structure 1PNF has been overlaid into the active sites of these models. (b) Genomic context of PNGaseRz with the putative proteins encoded by the genes annotated.

The genetic context of the genes encoding the PNGases was also considered, as genes encoding complementary functions are often grouped together, especially in bacterial genomes. Most of these genes were orphans, but the gene encoding PNGaseRz is likely part of an operon encoding putative enzymes with probable activity against N-glycans: α-fucosidases, an α-mannosidase, a glycoside hydrolase 18 family member, and an asparaginase (figure 3).

### Biochemical characterisation of the six PNGases

To investigate the activities of the putative PNGase enzymes, they were recombinantly expressed in *E. coli* (Supplementary figure 1) and assays were carried out with six glycoprotein substrates with varying types of N-glycans. The release of glycans was observed using thin layer chromatography and the decrease in protein size from N-glycan removal was monitored through SDS-PAGE (figure 4). The SDS-PAGE gels were stained for both total protein and glycoprotein. The substrates decorated with mammalian complex N-glycans (α1acid glycoprotein and fetuin) and high-mannose N-glycans (RNase B) were also treated with α-sialidase and α1,2-mannosidase, respectively, to provide simpler N-glycan structures to provide the best chance of observing activity. PNGaseL showed activity against all substrates tested, which included mammalian complex, high-mannose, plant-type and invertebrate-type N-glycans. The other five enzymes showed narrower specificity. The most common activities were observed against bovine RNase B and Honeybee venom phospholipase A2, which have high-mannose and invertebrate-type N-glycan decorations, respectively.

**Figure 4.**
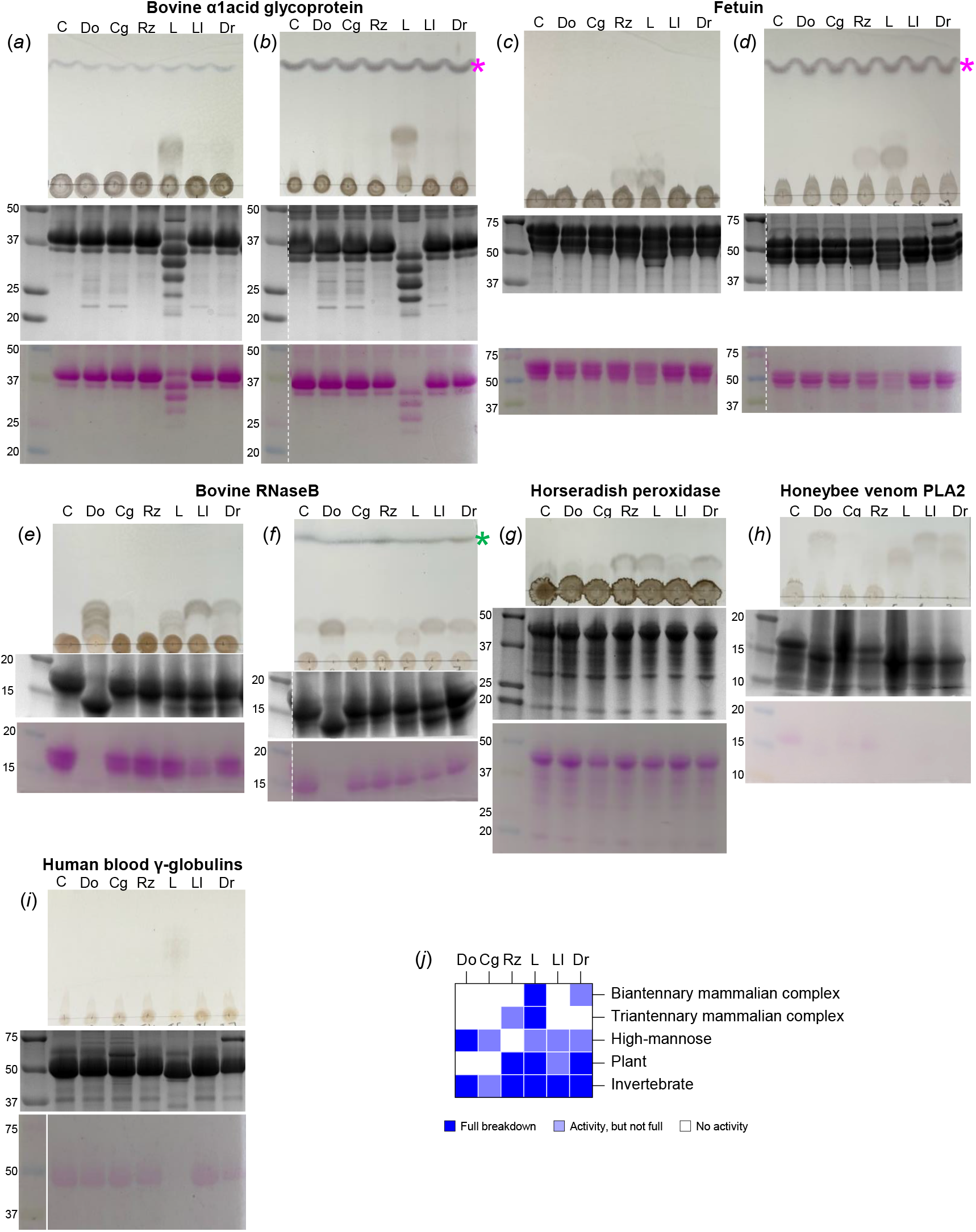
Initial specificity screening for activity of the six PNGase enzymes. Each panel has a TLC to visualise the glycan being released from the glycoprotein (top) and two SDS-PAGE gels to visualise the decrease in protein MW, stained for protein (middle) and glycoprotein (bottom). The PNGase tested is labelled with its two letter code and C – control. a) Bovine α1acid glycoprotein (biantennary mammalian complex N-glycans). b) Same as a) with the addition of a sialidase. c) Bovine fetuin (triantennary mammalian complex N-glycans). d) Same as c) with the addition of a sialidase. e) Bovine RNAseB (high-mannose N-glycans). f) Same as e) with the addition of an α1,2-mannosidase. g) Horseradish peroxidase (plant-type N-glycans, no antenna). h) Honeybee venom PhospholipaseA2 (insect-type N-glycans, no antenna). i) Human blood immunoglobulins (mix of mammalian N-glycans). j) A summary of the results. C – control, pink asterisk – indicates the sialic acid, green asterisk – indicates mannose.

### Investigating the structural basis for specificity displayed by the different PNGases

To investigate the specificity displayed by the six PNGases at a structural level we compared the active sites of the Alphafold models to four crystal structures – PNGaseF (1PNF), PNGaseF-II (4R4X, PNGaseBf (*Bacteroides fragilis*; 3KS7) and PNGasePm (*Phocaeicola massiliensis*; 7ZGN)^6,7,24^. PNGaseF and PNGaseBf have broad activity towards N-glycans without core α1,3-fucose decorations^7^. In contrast, PNGaseF-II and PNGasePm have specificity towards N-glycans with this core fucose (requirement for activity) and there is a pocket in the active site to accommodate this sugar^6,7^. To compare the active sites, we coloured the different loops extending out of the core β-sheets on the side of the protein with the active site and overlaid chitobiose from PNGaseF (figure 5). Active site residues were selected that likely interact with substrate based on previous literature (Supplementary figure 2) and a surface representation to visualise the contribution of the different loops to the active site (Supplementary figure 3).

**Figure 5.**
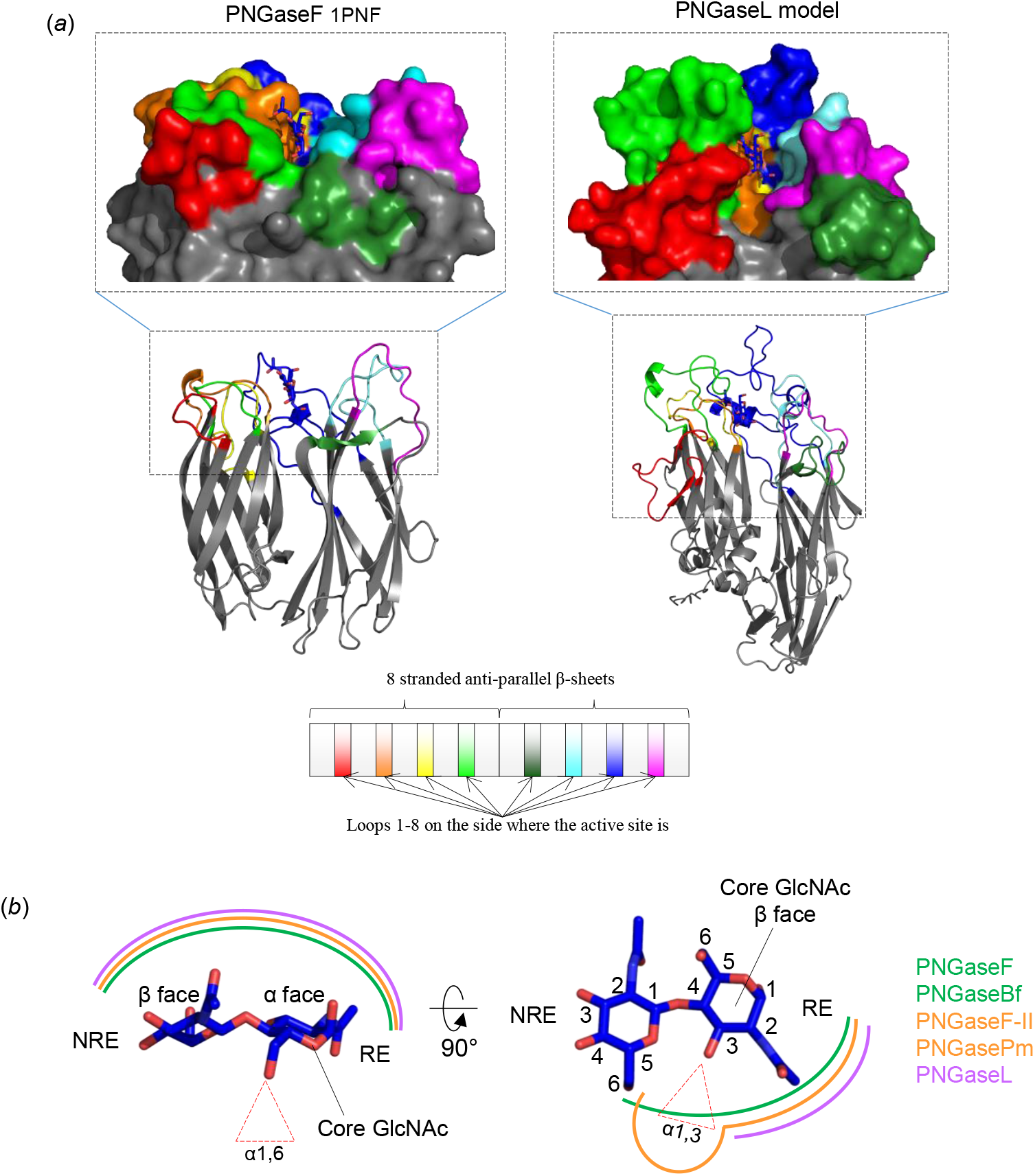
Structural comparison of PNGaseF structure and PNGaseL model. a) PNGaseF and L are shown as cartoons to emphasise their similarity regarding the structure of the catalytic module. The loops extending from the β-sheets have been coloured according to the diagram to indicate where the contributing interactions with the core N-glycan chitobiose originate. The loops are the most variable parts of the catalytic module between PNGase structures. The chitobiose from PNGaseF (1PNF) has been overlaid into the PNGaseL model to provide context. b) The interactions of different PNGases with the core N-glycan chitobiose can be categorised into three types indicated by different colours. Interactions with the α-face of the core GlcNAc and the β-face of the second GlcNAc is consistent between PNGases (left). The interactions around the C3 of the core N-glycan GlcNAc and C6 of the second GlcNAc vary (right). PNGaseF forms relatively close contact and prohibits the accommodation of an α1,3-fucose (green), but PNGaseF-II has a pocket conferring specificity towards this sugar (orange). In contrast, PNGaseL provides a much more open space in this area to allow broad specificity (purple). RE and NRE stand for reducing end and non-reducing end, respectively.

Comparison of the active site residues that likely interact directly with the substrate, reveal three broadly conserved features, which are provided by residues coming from the orange, lime green, and cyan loops in most cases (Supplementary figure 2). The orange loop contributes an aromatic-D-positive/aromatic motif. All three of these residues sit on the α-face of the first GlcNAc, but the first aromatic residue and third aromatic/positive also likely interact with the amide group from the second GlcNAc. The orientations of these aromatics are such that it is unlikely they provide hydrophobic platforms for the GlcNAc to stack against. The cyan loop predominantly interacts with the β-face side of the first GlcNAc through a histidine and glutamic acid. The exception to this is PNGaseF, which has a tryptophan instead of a histidine. PNGaseL and PNGaseDr have an additional serine and phenylalanine, respectively, provided by the cyan loop, that may make interactions with this GlcNAc at C6. Finally, the lime green loop provides a tryptophan or tyrosine on the β-face of the second GlcNAc. The orientation of these aromatics suggests this residue provides a hydrophobic platform for the β1,4-linked mannose of N-glycans also. Notably, for PNGaseL and PNGaseDr, these aromatic residues are provided by the red loop instead. In the case of PNGaseDr, the red loop takes this role away from the lime green loop by almost completely shrouding the lime green loop (Supplementary figure 3). In contrast, the lime-green loop of PNGaseL is not buried but instead forms a relatively large structure above the second GlcNAc. From the model, it is unclear how this loop may interact with substrate, and it is possible this loop may take on a different conformation in the presence of substrate. Overall, the α-face of the second GlcNAc in the N-glycan core is the most exposed part of the glycan.

One of the most striking aspects of the models of the PNGase enzymes explored in this report, is the open space underneath the chitobiose, which contrasts what has been seen in crystal structures of PNGase enzymes solved so far (Figure 5). PNGaseF, F-II, Bf, and Pm all hold the chitobiose more extensively in the region of C3 of the core GlcNAc and C6 of the second GlcNAc than the six new PNGase models analysed here. This complements the observed activities against substrates with α1,3-fucose on the core GlcNAc (plant and invertebrate) for the six PNGases described here. In contrast, the activity of PNGaseF-II and PNGasePm against substrates with core α1,3-fucose is facilitated by a pocket to accommodate this fucose rather than this area being completely open^6,7^.

One question that remains unanswered from the models is the reason why most of these PNGase enzymes cannot act on mammalian complex N-glycans, even the relatively simple de-sialylated biantennary structures found on α1acid glycoprotein. There are no obvious structural characteristics of these enzymes that would sterically hinder the accommodation of these structures compared to high-mannose N-glycans. However, no structures of PNGases have been solved with anything larger than chitobiose, which demonstrates there is still much to be understood about how structure relates to function for this family of enzymes.

### Comparison of PNGaseL activity to PNGase F and A

The broad specificity displayed by PNGaseL led us to compare its activity in more detail with the main PNGases used in glycobiology, PNGaseF and A. These three PNGases were tested against glycoprotein substrates decorated with a range of different N-glycan structures and prepared under three conditions: 1) untreated 2) boiled and 3) boiled and detergent (following the commercial PNGaseF protocol). The released glycans were labelled with procainamide and analysed by liquid chromatography-fluorescence detection-electrospray ionisation-mass spectrometry (LC-FLD-ESI-MS). The results are summarised as a heat map (figure 6) and full data is presented in Supplementary figure 4. PNGaseF was able to fully remove mammalian complex and high-mannose N-glycans from different glycoproteins when they had been pre-treated by boiling and detergent, but unable to remove plant- or invertebrate-type N-glycans (α1,3-core fucose decorated) in line with previous observations. This activity was slightly reduced under the boil-only conditions and vastly reduced under the untreated conditions. In contrast, PNGaseA was only able to act on plant N-glycans and the activity was not very high under any condition. This is likely due to PNGaseA preferring substrates where the protein has been partially digested^10^. PNGaseL was able to remove N-glycans from all the glycoproteins tested and, where there was overlap in specificities, the activity was largely comparable to PNGaseF. The results for honeybee venom phospholipase A demonstrate this (figure 6). It is of note that most of the N-glycans observed for honeybee venom phospholipase A do not have core fucosylation. This likely explains why activity against the insect N-glycans was common during the initial screening of the six PNGases (figure 4). From this screen, the data suggest that PNGaseL does remove N-glycans with bisecting GlcNAcs, but it does not do this as well as PNGaseF for some substrates.

**Figure 6.**
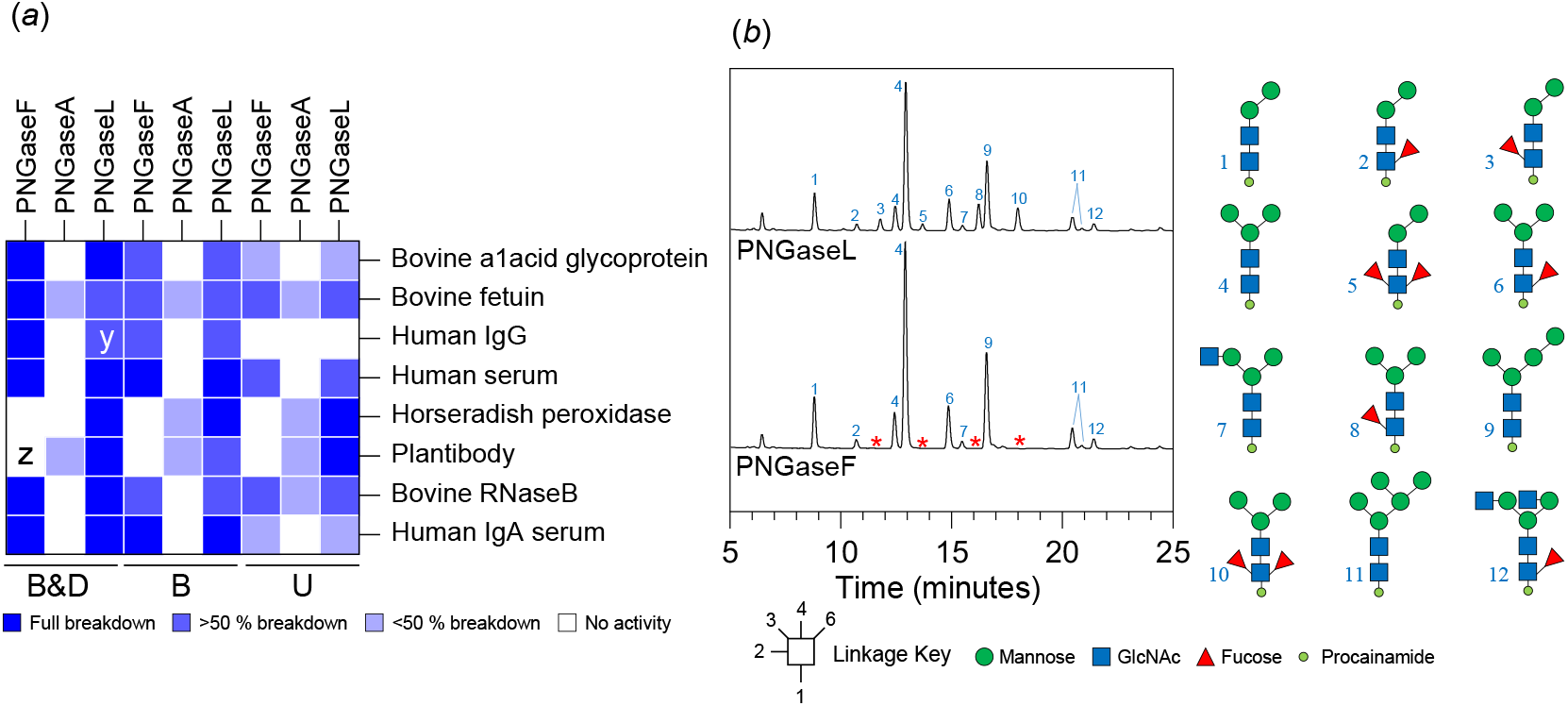
**Comparison** of PNGaseL activity to PNGaseF and A. (*a*) A heat map summary of the recombinant PNGaseL activity against a variety of N-glycosylated proteins in comparison to PNGaseF and PNGaseA The dark blue and white indicate full and no activity, respectively, and partial activities are represented by lighter blues. B&D, B and U represent ‘boiled and detergent’, ‘boiled’ and ‘untreated’, respectively. x:lower release of N-glycans with branched sialic acids compared to PNGaseF, y:low release of bisecting N-glycans, z:only high-mannose N-glycans released. Bovine α1acid glycoprotein, bovine fetuin, human IgG, human serum, horseradish peroxidase, plantibody and Bovine RNaseB have biantennary complex N-glycans, triantennary complex N-glycans, biantennary complex N-glycans with some bisecting GlcNAc and core α1,6 fucosylation, biantennary complex N-glycans with some core α1,6-fucosylation, simple plant N-glycans, variable plant N-glycans and high-mannose N-glycans, high-mannose N-glycans, and predominantly biantennary complex N-glycans with some core α1,6-fucosylation and bisecting GlcNAc, respectively. The full data is presented in Supplementary figure 4. (*b*) LC-ESI-FLD-MS data of N-glycans from honeybee venom phospholipaseA2 released by PNGaseL and PNGaseF. Peaks missing from the PNGaseF assay are indicated by a red asterisk and are all glycans with α1,3-core fucose.

### Assessing PNGaseL activity parameters and stability

A series of validation studies were carried out to find the optimal storage and assay conditions for PNGaseL and then to test its robustness and lifespan. In terms of stability, we carried out Differential Scanning Fluorimetery (DSF) using different buffers, pHs, and additives. We found that PNGaseL was most stable in 20 mM citrate buffer, 100 mM NaCl, at pH 6. Different pH was then assessed for optimal activity and assays carried out at pH 7.5 were found to release more glycan than pH 6 (Supplementary figure 5). This is the case for both bovine fetuin and human IgG. The stability of PNGaseL was then assessed by storing at different temperatures (room temperature and 37 °C) and over time (7 days and 14 days). Assays were then run against horseradish peroxidase and human IgG (only 7 days). The results showed that PNGaseL remained active under these parameters.

Finally, the ability of PNGaseL to remove N-glycans decorated with bisecting GlcNAc was explored in more depth using chicken egg white ovalbumin. The results confirm that PNGaseL releases these glycans, albeit less efficiently than PNGaseF (Supplementary figure 5). The active site of the PNGaseL model has a tyrosine from loop 1 (Y38) that is approximately in the same position as W200 from loop 4 of the PNGaseF crystal structure. When the chitobiose from the PNGaseF 1PNF structure is overlaid into the PNGaseL active site then this tyrosine is significantly closer to the C4 of the GlcNAc where the bisecting sugar would attach than the W200 is in the PNGaseF structure. The closer positioning of this tyrosine may sterically hinder N-glycans with bisecting GlcNAcs. However, direct evidence of this would need to be confirmed with structural data and this is only a logical suggestion for why PNGaseL struggles to accommodate this decoration.

## Discussion

PNGaseF is extensively used in glycobiology to allow release of the N-glycome of the target protein, cell or tissue under study. While the enzyme has a wide specificity for N-glycans, it requires certain conditions for optimal activity and is unable to accommodate core α1,3-fucose decorations found in plant and invertebrate N-glycans. In this study, six PNGase enzymes were characterised from sources that are not associated with mammals. Overall, activity against high-mannose N-glycans was the most common among these six and this is reflective of these glycans having a consistent structure across kingdoms. Screening against substrates with different N-glycans revealed some notable observations. For instance, PNGaseRz selects specifically for α1,3 core-fucosylated glycans and this has been observed previously for PNGaseFII^6^. There was not much activity observed for PNGaseCg and it is likely that its true substrates have not been tested here as there is evidence of unusual N-glycan structures in oysters^19^.

PNGaseL was selected for more in-depth characterisation given its broad specificity for core fucose decorations. It is a robust enzyme with high potential for commercialisation and can be used under the same conditions as PNGaseF. The only shortcoming is the lower activity towards N-glycans with bisecting GlcNAcs compared to PNGaseF.

## Contributions

LIC, PAU, DIRS and DNB designed the research. CRB, LIC, PAU, CBT, JMMD and TOO performed research. MK and LJH carried out the bioinformatics. CRB and LIC prepared and revised manuscript. All authors contributed to editing the manuscript.

## Acknowledgements

The work was funded by The Academy of Medical Sciences (SBF0061175), the Wellcome Trust and Royal Society Sir Henry Dale fellowship (224240/Z/21/Z) awarded to L.I.C. and a BBSRC/Innovate UK IB catalyst award to D.N.B. ‘Glycoenzymes for Bioindustries’ (BB/M029018/1). T.O.O. is funded by the BBSRC Midlands Integrative Biosciences Training Partnership (MIBTP) with his studentship in collaboration with industrial partners Ludger (Oxford, UK) awarded to L.I.C.

---

**Supplementary figure 1.**
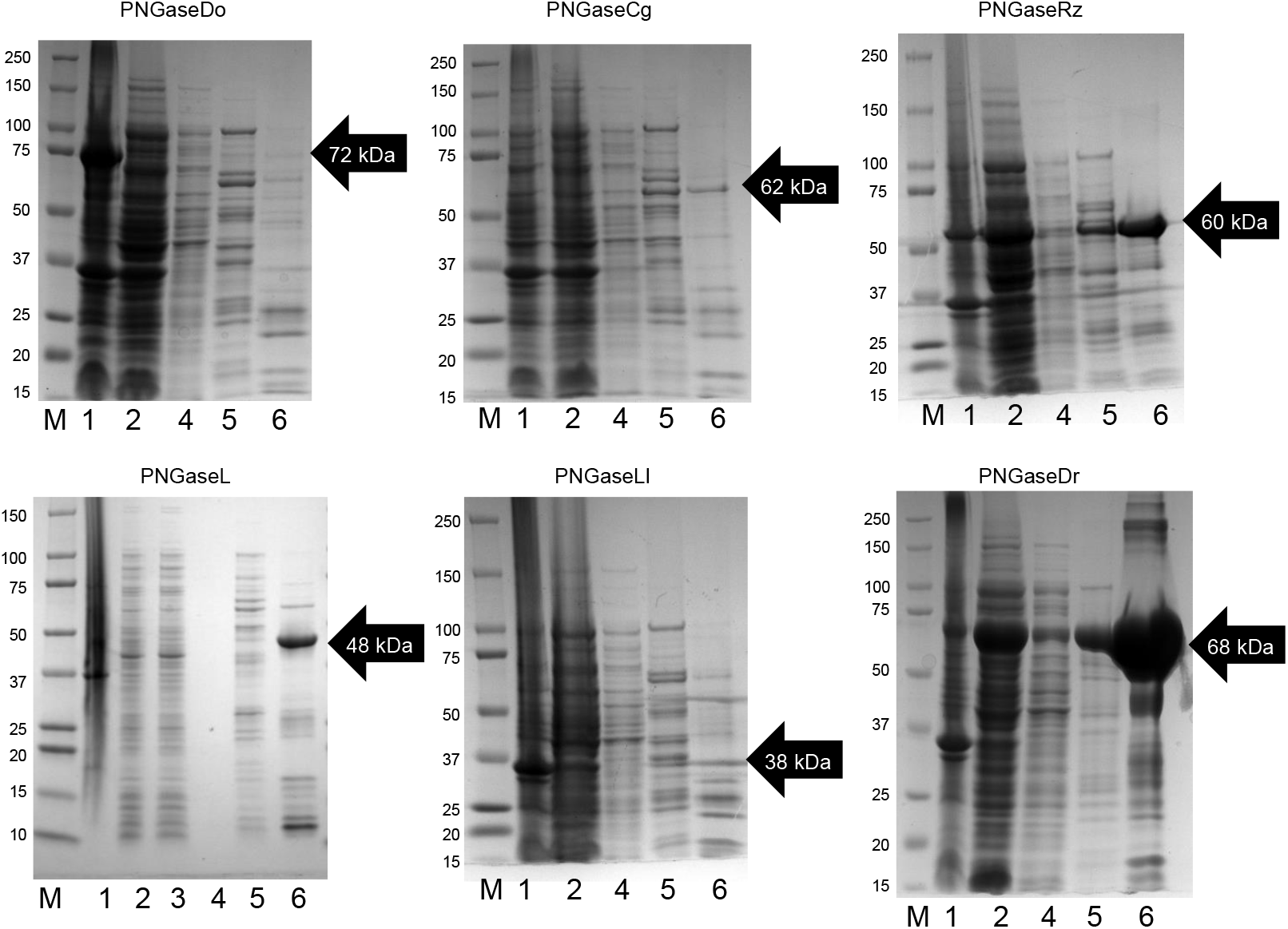
Purification of six putative recombinant PNGase enzymes. SDS-PAGE gels of the purification steps for the PNGase enzyme. This is a relatively crude initial purification to assess the success of recombinant expression. 1-6 is Pellet, cell lysate, column flow-through, buffer wash, 10 mM imidazole wash and 100 mM imidazole. M – marker and the bands are labelled with kDa of the standard proteins.

**Supplementary figure 2.**
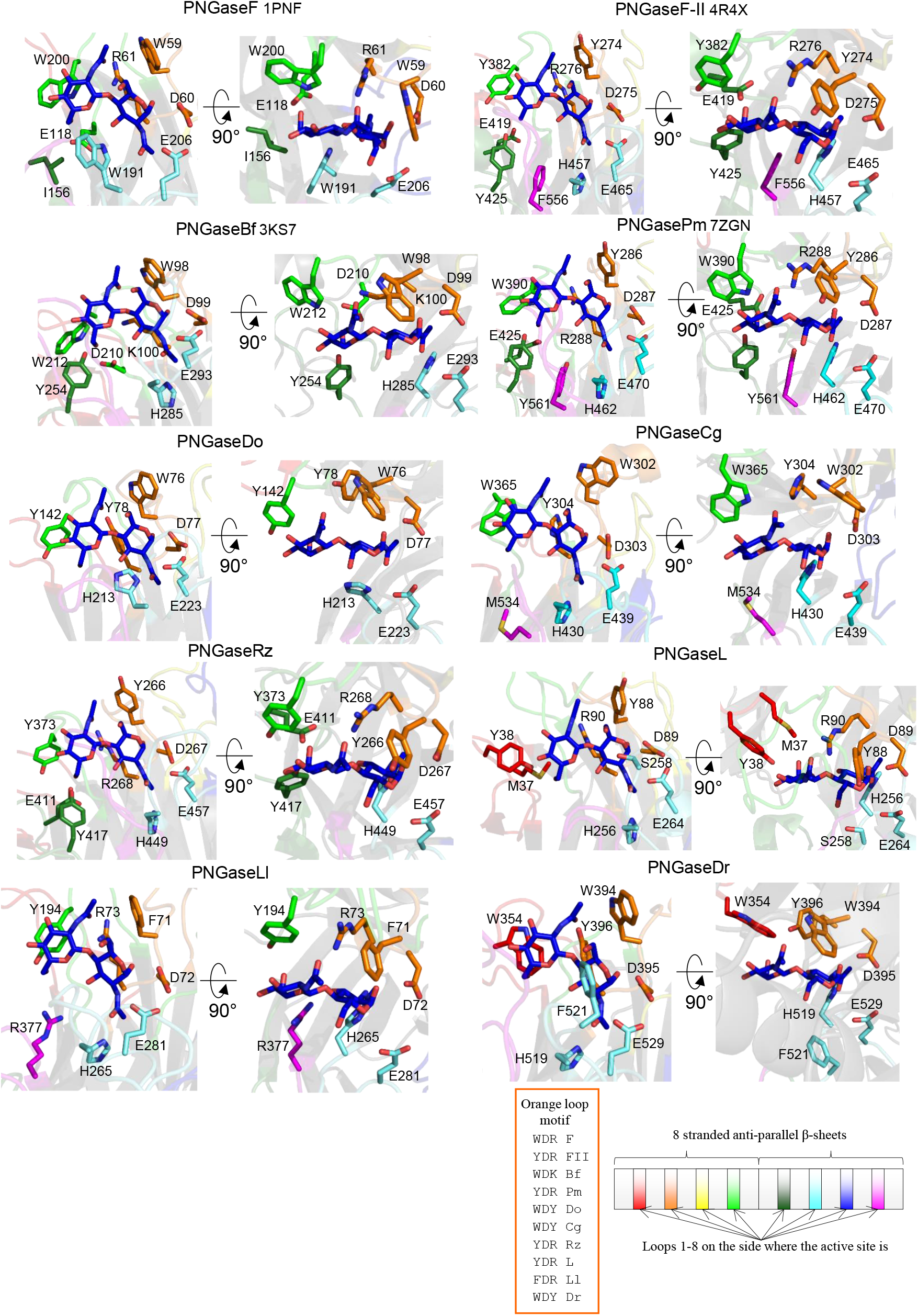
The active sites of PNGase crystal structures and models. The loops have been coloured according to the diagram (right) to indicate where the contributing residues come from. The chitobiose from PNGaseF (1PNF) has been overlaid into each structure/model to provide context and the residues likely contacting the chitobiose are shown as sticks.

**Supplementary figure 3.**
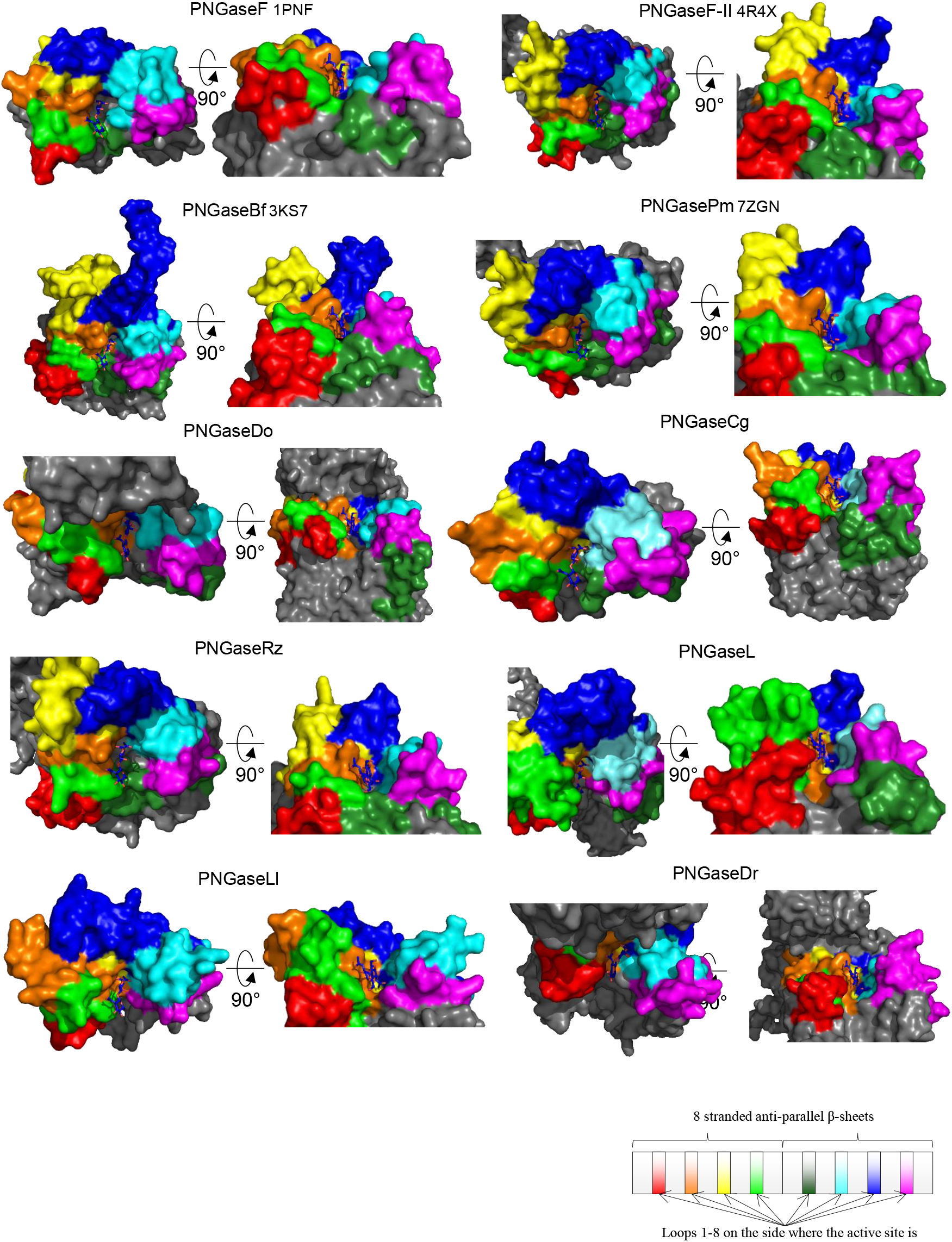
**Surface representation of the** The contribution of different loops to the active sites of PNGase enzymes. The loops have been coloured according to the diagram (right) to indicate where the contributing residues come from. The chitobiose from PNGaseF (1PNF) has been overlaid into each structure/model to provide context. Each enzyme is shown from two angles – one from above and one looking into the active site at the non-reducing end of the chitobiose.

**Supplementary figure 4.**
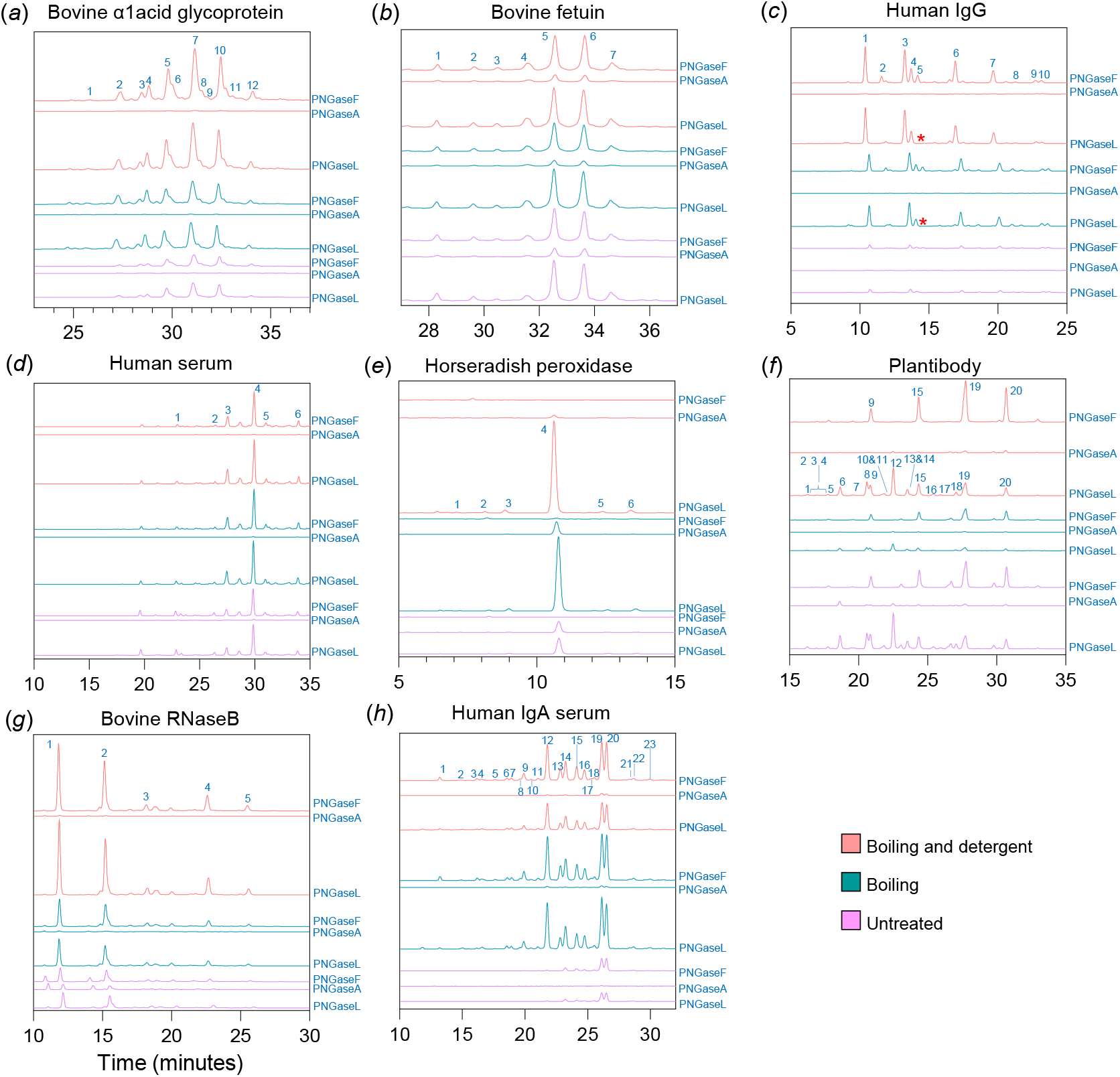
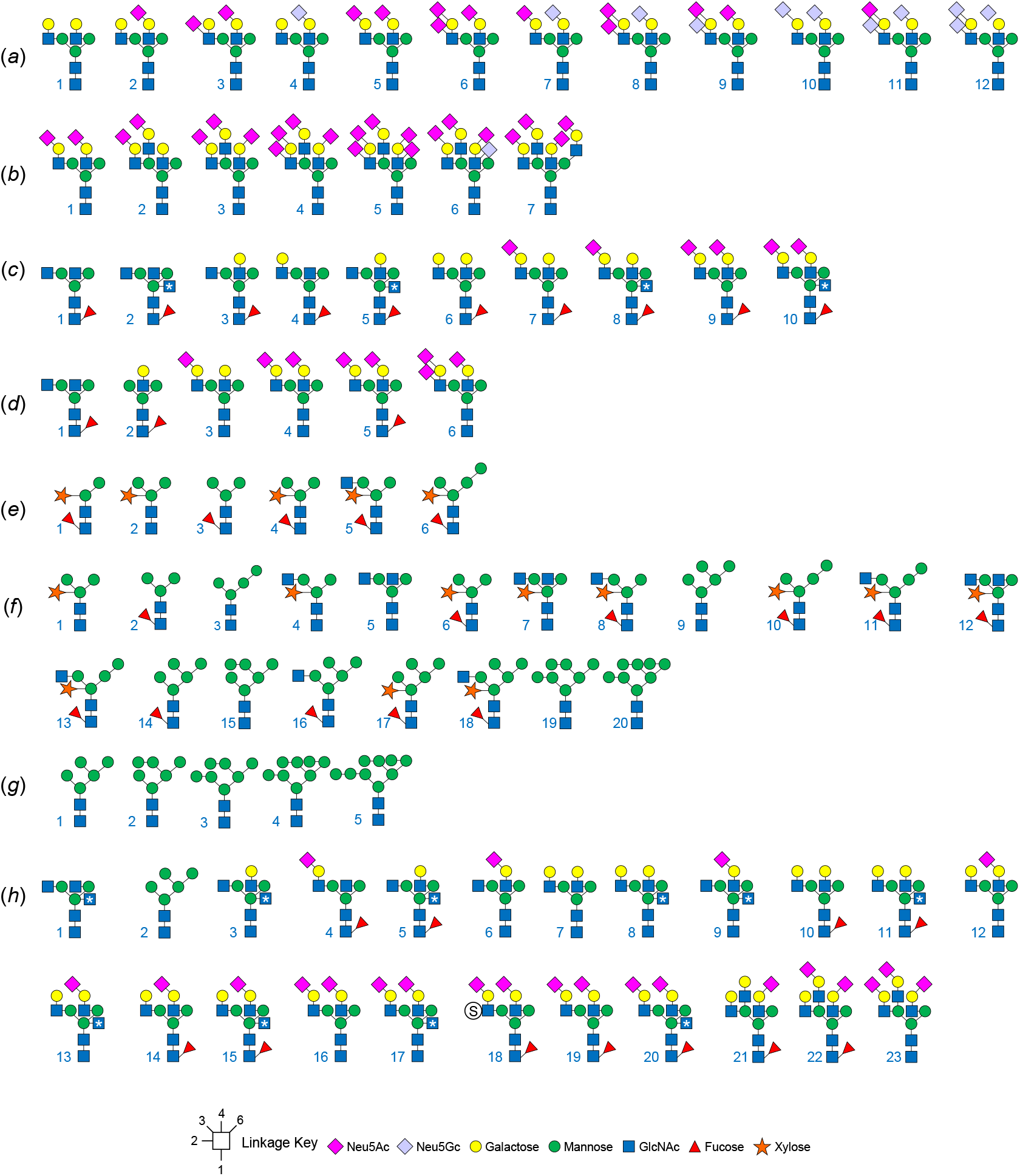
Activity of three PNGases against glycoprotein substrates under three different conditions. Each panel is a different glycoprotein substrate, the colours of the chromatograms correspond to different assay conditions and the peaks are numbered to indicate the different glycan structures (next page). Red asterisk indicate missing or reduced peaks.

**Supplementary figure 5.**
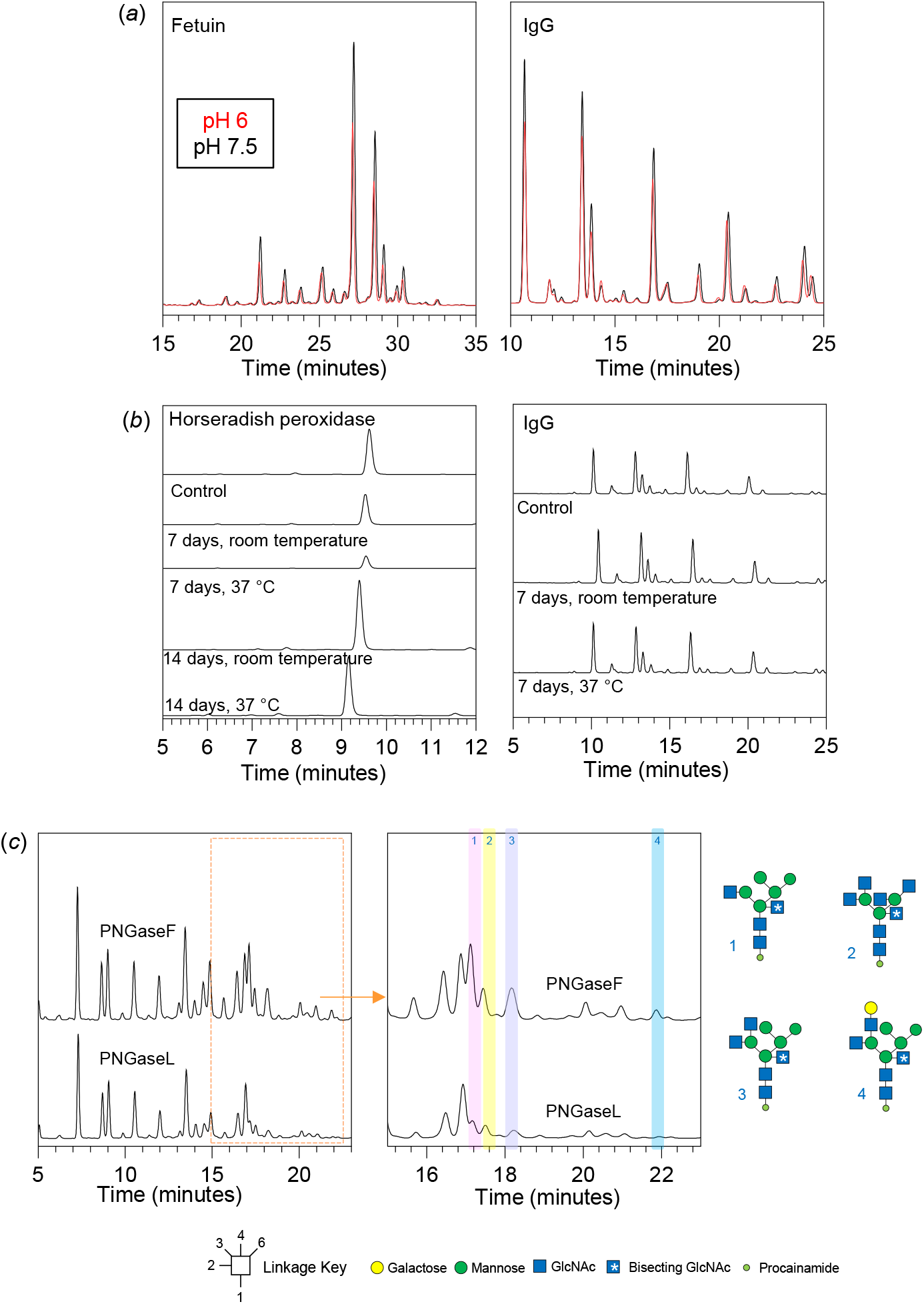
Assessing the different assay parameters for PNGaseL. (*a*) Assessing activity at different pH. (*b*) assessing stability of PNGase by storing it at different temperatures. (*c*) Assessing the ability of PNGaseL to remove bisecting N-glycans in more detail using chicken egg white ovalbumin. The left panel are the full chromatograms, and the right panel is the section where the bisecting N-glycans elute (orange box in left panel).

